# Efficient Isolation and Phenotypic Characterization of Primary Rat Pulmonary Pericytes

**DOI:** 10.64898/2026.07.27.740972

**Authors:** Dinesh Sundaram, Stuti Agarwal, Mathews V Varghese, Maki Niihori, Morgan L Nelson, Dinesh Bharti, Takanori Sano, Vinicio de Jesus Perez, Olga Rafikova, Ruslan Rafikov, Joel James

**Affiliations:** Division of Pulmonary, Critical Care, Sleep and Occupational Medicine, Department of Medicine, Indiana University, Indianapolis, Indiana; Stanford School of Medicine, Stanford University, Stanford, California

## Abstract

Pericytes are essential regulators of pulmonary vascular homeostasis, but their isolation from rat lung tissue is challenging because no single marker uniquely identifies them and contaminating fibroblasts, endothelial cells, hematopoietic cells, and vascular smooth muscle cells can persist during isolation and culture. Here, we describe a rapid and efficient protocol for the isolation of primary rat pulmonary pericytes using sequential magnetic depletion of CD45-positive hematopoietic cells and CD31-positive endothelial cells, followed by positive selection for NG2-positive cells. The isolated cells were expanded in culture and characterized by immunofluorescence and functional co-culture assays. Cultured cells displayed typical pericyte morphology and expressed the pericyte-associated markers NG2, PDGFRβ, and 3G5, with minimal expression of CD31, CD45, PDGFRα and MYH11. In endothelial cell-pericyte ECM gel co-culture assays, isolated pericytes associated with endothelial cords and were frequently observed near network branch points and junctions, further supporting their pericyte identity. This method yields an enriched population of primary pulmonary pericytes suitable for downstream applications, including cell culture and functional studies. Overall, this streamlined protocol provides a practical platform for studying pulmonary pericyte biology in rat models of health and vascular disease.

---

Pulmonary vascular diseases are characterized by progressive alterations in the structure and function of the pulmonary circulation^1^. Rat models are widely used to investigate pulmonary vascular remodeling, right ventricular adaptation, disease progression, and responses to potential therapies^2^. The utility of these models depends in part on reliable methods for isolating the individual pulmonary vascular cell populations that contribute to disease. In particular, isolation of primary pulmonary pericytes is important for defining pericyte-specific mechanisms and developing therapeutic approaches directed at restoring microvascular function.

Pericytes are mural cells closely associated with endothelial cells in capillaries and small vessels^3^. They contribute to endothelial survival, vascular stability, barrier integrity, angiogenesis, and microvascular remodeling^4,5^. Pericytes are also implicated in the pathogenesis of vascular diseases, including pulmonary hypertension, underscoring the importance of defining their roles in vascular homeostasis and remodeling^6^. However, their isolation from lung tissue is challenging because the lung contains multiple endothelial, hematopoietic, fibroblast, and smooth muscle cell populations with overlapping morphological and molecular features. Moreover, no single marker exclusively identifies all pericytes, making it necessary to combine positive enrichment with exclusion of contaminating cell types and validation using multiple phenotypic and functional criteria^7^.

The 3G5 antibody has long been used for the identification and isolation of pulmonary vascular pericytes^7-10^. However, 3G5 is not readily available commercially and generally must be produced from a hybridoma and subsequently purified, limiting its routine use^11^. In this study, we also describe the production and purification of 3G5 IgM from an established hybridoma for pericyte characterization. Neuron-Glial 2 (NG2) has been widely used as an alternative surface marker for isolating pericytes from human and mouse tissues, although its expression is not restricted exclusively to pericytes^8,12^. However, NG2-based enrichment can be effectively leveraged when combined with appropriate depletion strategies and selective culture conditions. Here, we describe a sequential magnetic-selection approach for isolating primary rat pulmonary pericytes through depletion of CD45-positive hematopoietic cells and CD31-positive endothelial cells, followed by enrichment of NG2-expressing cells. The isolated population was characterized by morphology, immunofluorescence, and endothelial cell-pericyte co-culture. This approach provides a practical method for obtaining enriched primary rat pulmonary pericytes for studies of pulmonary vascular biology, disease mechanisms, and therapeutic development.

## Methods

### Animals

All animal procedures were approved by the Indiana University Institutional Animal Care and Use Committee and were conducted in accordance with institutional guidelines for the care and use of laboratory animals. Male and female Sprague-Dawley rats, aged 10-13 weeks, were used for pulmonary pericyte isolation. Animals were maintained under a 12-hour light-dark cycle with free access to standard rodent chow and water. Rats were euthanized using an institutionally approved method, and the lungs were perfused with saline and collected immediately for cell isolation. At least three independent pericyte isolations were performed using lung tissue obtained from separate animals.

#### Isolation of primary lung pericytes

A detailed isolation protocol has been described below.

### Pericyte culture

Isolated cells were plated in a 10cm dish (Uncoated) and maintained in complete pericyte medium (ScienCell 1201). Cells were cultured at 37°C in a humidified incubator containing 5% CO_2_. Medium was replaced every 2 days, and cells were passaged at approximately (70-90%) confluence with trypsin-EDTA. Cells between passages 1 and 2 were used for characterization and subsequent experiments.

### Immunofluorescence staining

Primary lung pericytes were cultured on glass coverslips and fixed with 4% paraformaldehyde for 15 minutes at room temperature. Cells were washed with phosphate-buffered saline and, where required for intracellular antigens, permeabilized with 0.1% Triton X-100. Nonspecific binding was blocked with 10% bovine serum albumin for 30 minutes at room temperature. Cells were incubated with primary antibodies against CD31/PECAM1 (Novus AF3628), CD45 (Invitrogen, MA1-81566), PDGFRα (Novus AF1062), MYH11 (Novus NBP2-44533) NG2/CSPG4 (Sigma-Aldrich AB5320), PDGFRβ (Novus NBP2-46361), or purified 3G5 IgM. Unless otherwise indicated, primary antibodies were used at a dilution of 1:100. Purified 3G5 IgM was used at 2 µg/mL. Rabbit primary antibodies were detected using chicken anti-rabbit IgG conjugated to Alexa Fluor 488 (Invitrogen, catalog no. A-21441; 1:250), and 3G5 IgM was detected using goat anti-mouse IgM conjugated to DyLight 650 (Invitrogen, SA5-10153; 1:500). Cells were mounted with ProLong™ Diamond Antifade Mountant with DAPI (Invitrogen P36962) and imaged using a Zeiss Axio Observer 7 or ECHO Revolve microscope. Image-acquisition settings were maintained consistently between relevant samples and controls.

### Preparation of 3G5 IgM from hybridoma supernatant

The 3G5 hybridoma cell line (ATCC CRL-1814) was cultured in ATCC-formulated Dulbecco’s modified Eagle’s medium supplemented with 10% fetal bovine serum at 37°C and 5% CO2 as previously described^10,13^. The hybridoma produces a mouse IgM monoclonal antibody recognizing the 3G5 antigen expressed on microvascular pericyte plasma membranes. Approximately 100 mL of hybridoma culture supernatant was clarified by centrifugation at 4,000 × g for 10 minutes and filtered through a 0.22-µm membrane. The supernatant was concentrated using Amicon Ultra-15 centrifugal filters and buffer-exchanged into ice-cold binding buffer containing 20 mM Tris and 1.25 M NaCl, pH 7.4. IgM was purified using the Pierce IgM Purification Kit (Thermo Fisher Scientific 44897). The concentrated sample was mixed 1:1 with binding buffer, applied to the equilibrated column, incubated for 30 minutes at 4°C, washed, and eluted following a 1-hour incubation with elution buffer at room temperature. The purified IgM was concentrated and buffer-exchanged into sterile PBS using Amicon centrifugal filters. Antibody concentration was measured using a NanoDrop spectrophotometer. Glycerol was added to a final concentration of 5%, and the antibody was aliquoted and stored at -80°C. Antibody integrity was assessed by SDS-PAGE based on the expected IgM heavy- and light-chain bands.

### Endothelial cell-pericyte ECM gel co-culture assay

Following a modified protocol as described^13^, ECM gel derived from Engelbreth-Holm-Swarm murine sarcoma, growth factor-reduced and phenol red-free (Sigma-Aldrich, catalog no. E6909-5ML), was thawed overnight on ice. The gel was added to a 96-well plate at 55 μL per well and allowed to polymerize for 30 minutes at 37°C. Pulmonary microvascular endothelial cells were labeled with the CellTrace CFSE Cell Proliferation Kit (Thermo Fisher Scientific, catalog no. C34570), and primary rat pulmonary pericytes were labeled with the CellTrace Far Red Cell Proliferation Kit (Thermo Fisher Scientific, catalog no. C34572), according to the manufacturer’s instructions. Approximately 5,000 endothelial cells and 1,000 pericytes were combined in 100 μL of pericyte medium containing 2% FBS and seeded onto each ECM gel-coated well. After 4 hours of co-culture at 37°C, fluorescence images were acquired using a 4X objective on an ECHO Revolve microscope.

## Results

### Isolation and characterization of primary lung pericytes

Lung tissue was finely minced and enzymatically dissociated with collagenase for 45 minutes at 37°C (Fig. 1A). The resulting cell suspension was filtered and washed before depletion of hematopoietic and endothelial cells using antibodies against CD45 and PECAM1/CD31. The CD45^−^/PECAM1^−^ fraction was subsequently subjected to positive selection for NG2, and the NG2-positive cells were collected and cultured in pericyte growth medium. Following attachment and expansion, the isolated cells displayed an elongated, stellate morphology with multiple cellular processes, consistent with the morphology of cultured pericytes (Fig. 1B). Immunofluorescence staining demonstrated prominent expression of the pericyte-associated markers NG2 and PDGFRβ. In contrast, CD31 staining was minimal, indicating limited contamination by endothelial cells (Fig. 1B).

**Figure 1.**
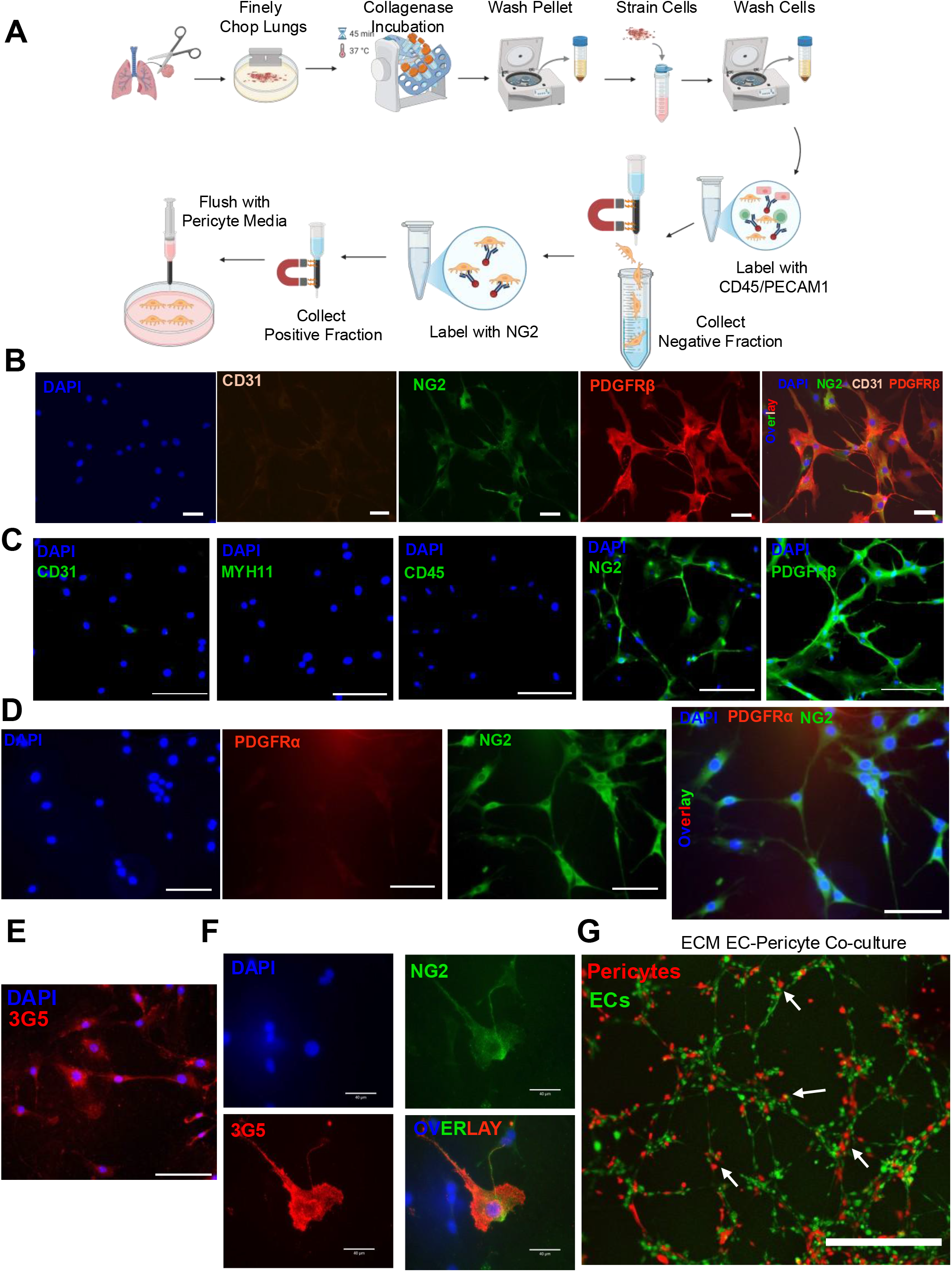
Isolation and characterization of primary lung pericytes. **A**, Schematic representation of the lung pericyte isolation procedure. Lungs were finely minced, digested with collagenase for 45 minutes at 37°C, washed, and passed through a cell strainer. CD45- and PECAM1/CD31-positive cells were removed by negative magnetic selection. The negative fraction was subsequently labeled for NG2, and the NG2-positive fraction was collected and cultured in pericyte medium. **B**, Representative immunofluorescence images of cultured cells co-stained for DAPI, CD31, NG2, and PDGFRβ, with a merged image showing expression of the pericyte markers NG2 and PDGFRβ. Scale bars: 50μm **C**, Representative images demonstrating minimal staining for CD31, CD45, and MYH11 and prominent staining for NG2 and PDGFRβ. Scale bars:100μm **D**, Representative images demonstrating minimal staining for fibroblast marker PDGFRα and positive for NG2. Scale bars 90μm **E**, Representative staining of isolated cells for the pericyte-associated marker 3G5. Scale bars: 100μm **F**, Representative images showing co-expression of NG2 and 3G5. Scale bars:40μm. **G**, Functional association of isolated pulmonary pericytes with endothelial networks. Pulmonary microvascular endothelial cells isolated as recently described^14^ were labeled with CellTrace CFSE are shown in green, and primary rat pulmonary pericytes labeled with CellTrace Far Red are shown in red. Cells were co-cultured on growth factor-reduced, phenol red-free ECM gel derived from Engelbreth-Holm-Swarm murine sarcoma. Pericytes were observed along endothelial cords and near network branch points and junctions. White arrows indicate representative sites of endothelial-pericyte association. Scale bar, 500μm. The images are representatives of three independent experiments.

Further phenotypic characterization showed little to no detectable expression of CD31, CD45, or the smooth muscle cell marker MYH11, whereas the cultures exhibited strong staining for NG2 and PDGFRβ (Fig. 1C). Minimal PDGFRα staining was observed, consistent with limited contamination by PDGFRα-expressing fibroblast-like cells (Fig. 1D). Furthermore, the isolated cells also expressed the pericyte-associated surface antigen 3G5 (Fig. 1E). Co-immunofluorescence analysis demonstrated that 3G5-positive cells co-expressed NG2 (Fig. 1F). Across all isolations, cultured cells consistently displayed pericyte-like morphology, prominent NG2, PDGFRβ, and 3G5 expression. To evaluate the ability of the isolated pulmonary pericytes to interact with endothelial vascular structures, fluorescently labeled pericytes were co-cultured with pulmonary microvascular endothelial cells on growth factor-reduced, phenol red-free ECM gel. Endothelial cells formed interconnected tube-like networks, while pericytes associated with the endothelial cords and were frequently observed near branch points and junctional regions of the network (Fig. 1G). This pattern provides functional support for the pericyte identity of the isolated cells. Together, these findings support the successful enrichment of a primary lung pericyte population with minimal detectable endothelial, hematopoietic, fibroblast or mature smooth muscle cell contamination.

## Discussion

In this study, we demonstrate a sequential magnetic-selection approach for isolating primary rat pulmonary pericytes. The method combines depletion of CD31-positive endothelial cells and CD45-positive hematopoietic cells with enrichment of NG2-expressing cells. The isolated cells displayed the elongated, stellate morphology typical of cultured pericytes, expressed NG2, PDGFRβ, and 3G5, and showed minimal detectable expression of CD31, CD45, the fibroblast-associated marker PDGFRα, and the mature smooth muscle cell marker MYH11. Their association with endothelial cords and network junctions in ECM gel co-culture provided additional functional support for their pericyte identity^13,15^. A major strength of this approach is the use of sequential depletion and enrichment rather than reliance on a single marker. Although NG2 is widely used for pericyte isolation, it is not exclusively expressed by pericytes and may also be present in other mural or mesenchymal populations. Depletion of CD31-positive and CD45-positive cells before NG2 selection reduces major sources of endothelial and hematopoietic contamination. Nevertheless, the isolated population should be validated using a combination of pericyte-associated markers, exclusion markers, morphology, and functional interaction with endothelial cells^13,15^.

Post-isolation culture conditions were also important for maintaining an enriched pericyte population. Frequent medium changes during early colony formation helped remove debris, dead cells, and poorly adherent contaminants. Cells were maintained on uncoated culture surfaces because we found that gelatin coating promoted the expansion of cells with fibroblast-like morphology during protocol optimization. Low-serum pericyte medium provided an additional selective environment by supporting pericyte growth while limiting the expansion of fibroblasts and vascular smooth muscle cells, which generally proliferate more readily under higher-serum conditions. These effects represent practical observations from protocol development rather than formal comparisons of culture conditions. When fibroblast-like cells persist, a brief CD90-depletion step may be used to remove cells with high CD90 expression and further enrich the pericyte population. This step should be applied cautiously, because pericytes themselves can express CD90, and both CD90-positive and CD90-negative pericyte populations have been described^16^. Therefore, CD90 depletion is best reserved for cultures with clear fibroblast-like overgrowth rather than used routinely. Because prolonged culture may promote phenotypic drift, early-passage cells are recommended, and marker expression should be reassessed when later-passage cells are used. The cells could generally be expanded through approximately passages 5-6, although early-passage cells are recommended and later-passage cultures should be revalidated for marker expression and morphology. A single round of magnetic purification was typically sufficient; however, cultures with persistent contaminants may be expanded and subjected to repeat CD31/CD45 depletion, optional CD90 depletion, and NG2 reselection. Lung tissue from one rat was sufficient for routine isolation. When lungs from multiple animals are pooled, antibody and MicroBead volumes and column capacity should be scaled according to the total cell number.

A limitation of this protocol is that it may preferentially enrich an NG2-expressing pericyte subset and may therefore not capture the full heterogeneity of pulmonary pericytes^5^. In addition, the current characterization is largely qualitative. Flow cytometry would permit quantitative assessment of pericyte-associated and exclusion markers. Adaptation to human lung tissue will also require optimization because tissue quality, fibrosis, inflammation, and digestion efficiency may differ from rat lungs. Despite these limitations, the combination of endothelial and hematopoietic cell depletion, NG2-positive selection, selective culture conditions, and optional CD90 depletion provides a practical strategy for obtaining enriched primary rat pulmonary pericytes for mechanistic and therapeutic studies of pulmonary vascular disease.

## DETAILED PERICYTE ISOLATION PROTOCOL

### MATERIALS

- Basal DMEM + 1% antimycotic-antibiotic
- Complete DMEM (10% FBS + 1% antimycotic-antibiotic)
- Assay buffer: 0.5% biotin-free BSA in DPBS
- Fetal Bovine Serum (FBS)
- DNase I (10 mg/mL stock) (Sigma 11284932001)
- Collagenase Type IV (1 mg/mL final) (LS004188 Worthington)
- Neutral protease (1 mg/mL final) (Worthington LS02109)
- Endothelial Cell Isolation Kit (species-specific: Rat Miltenyi-130-109-679)
- CD45 MicroBeads, Rat- (Miltenyi, 130-109-682)
- Anti Rat NG2 antibody-Biotin conjugated (Sigma AB5320B)
- Streptavidin MicroBeads (Miltenyi, 130-048-102)
- LD (130-042-901) and LS columns (Miltenyi 130-042-401)
- MidiMACS separator
- 37°C water bath
- Refrigerated centrifuge (4°C, swinging bucket rotor)
- Cell strainers (70 µm, 40 µm, optional 30 µm)
- Gentle rotator

#### Digestion mix (prepare fresh)-10mL/1gm tissue

- Collagenase: 1 mg/mL
- Neutral protease: 1 mg/mL
- DNase I: 150 µL (10 mg/mL stock) per 10 mL
- Basal DMEM or DPBS

#### CRITICAL STEP

*Filter sterilize and keep on ice until use. Prior to use, warm to* 37°C.

##### Tissue Digestion

1. Add 0.5-1 g of the excised lung piece into a dish containing 500 µL of digestion mix.
2. Mince finely using a blade (∼ < 1 mm pieces) Transfer to 15 mL tube and adjust volume to 5-10 mL per 0.5-1 g tissue.
3. Incubate at 37°C for 45 min with gentle rotation.

#### CRITICAL STEP

*Avoid over digestion to preserve viability*.

4. Neutralize digestion with equal volume complete DMEM.
5. Pipette vigorously to dissociate clumps with 5 mL pipette.
6. Filter sequentially through 70 µm, then 40 µm strainers.
7. Centrifuge (400 × g, 10 min, 4°C).
8. Wash sequentially:
  - Basal DMEM (no FBS) once
  - Assay buffer twice
9. Resuspend and filter again (40 µm).
10. Adjust concentration to ≤1 × 10^7^ cells per 80 µL Assay buffer (∼500mg of tissue will yield this).

#### CRITICAL STEP

*Avoid over-loading cells for the depletion/selection steps*.

##### Labelling of Cells

###### Negative selection

1. Add 20 µL CD31 Antibody (from the Miltenyi EC isolation kit) and 20 µL CD45 MicroBeads.
2. Adjust volume to 200 µL with Assay buffer.
3. Incubate for 15 min at 4°C (**<u>not</u>** on ice).
4. Prepare LD column (rinse with 2 mL Assay buffer).
5. Apply cells to the LD Column (bring volume to 500 µL with Assay Buffer).
6. Collect flow-through in tube on ice (pericyte-enriched fraction).
7. Wash column 3X (1 mL each), collecting eluate.

###### Positive selection

8. Centrifuge eluate (400 × g, 5 min) and pool pellets.
9. Add 20 µL of Anti-Rat NG2-Biotin and 20 µL of Streptavidin Microbeads.
10. Incubate for 30-40 min at 4°C with gentle rotation.
11. Prepare LS column (rinse with 2 mL Assay buffer).
12. Apply cells and wash column 3 times with Assay buffer.
13. Discard flow-through.
14. Remove column from magnet.
15. Elute Pericytes with 5 mL Pericyte medium. Remove column from the separator and place it on a 15ml tube. Pipette 5 mL of Pericyte cell Media onto the column. Immediately flush out the magnetically labeled cells by firmly pushing the plunger into the column.
16. Plate the cells in a 6 well plate.
17. Change media on Day 3 post seeding and alternate days thereafter.
18. Colonies begin to form on day 4. Once sufficient colonies are formed, trypsinize and reseed into plate with same surface area.

#### CRITICAL STEP

*From day 3, change media every alternate day until adequate number of colonies form. Remove media gently when changing media (Use a pipette). Do not aspirate media completely when changing media*.

### NOTES

- Use biotin-free BSA in Assay buffer.
- Incubate cells at 4°C (not on ice) (antibody selection).
- Use Multiple LD/LS Columns if needed.
- Do not overload columns or overload cells during antibody tagging.
- Only add buffer when column reservoir is empty.

**Table 1.** Troubleshooting steps.

| Problem | Probable Cause | Solution |
| --- | --- | --- |
| Low Viability | Over digestion | Reduce incubation times and overall processing times. Collect cells on ice. |
| Clumping, filtration difficulty | DNA interference | Optimize DNase Concentration |
| Poor labelling | FBS interference | Remove FBS prior to labeling, Use biotin-free BSA |
| Contaminating Cells in Culture | Insufficient CD45 and CD31 antibody added | Optimize CD45 and CD31 antibody concentration<br>Use LD Column in depletion step instead of LS Column (Miltenyi -130-042-901) |
| Contaminating Fibroblast Cells in Culture | Media Changes are not done on alternate days. Use of Gelatin to Coat Plates. | Media Changes to be done on alternate days. Use CD90 microbeads (Miltenyi 130-121-273) to deplete fibroblasts and re-culture cells if needed (10-15 min incubation in 4°C) |

## Acknowledgments

We thank Dr. Ke Yuan for providing an aliquot of 3G5 antibody used during initial assay optimization. This work was supported by NIH grants R00HL171869 (JJ), R01HL151447, R01HL132918 (RR), R01HL133085, R01HL160666 (OR), R01HL139664, R01HL134776, R01HL59886, R01HL160018, 5R01HL172449-02 (VDJ) and AHA grant 23CDA1050843 (MN). Elements of figure 1A were created using Biorender.

## References

1. Ejikeme C, Safdar Z. Exploring the pathogenesis of pulmonary vascular disease. Front Med (Lausanne). 2024;11:1402639.

2. Wu XH, Ma JL, Ding D, Ma YJ, Wei YP, Jing ZC. Experimental animal models of pulmonary hypertension: Development and challenges. Animal Model Exp Med. 2022;5(3):207–216.

3. Bergers G, Song S. The role of pericytes in blood-vessel formation and maintenance. Neuro Oncol. 2005;7(4):452–464.

4. Payne LB, Hoque M, Houk C, Darden J, Chappell JC. Pericytes in Vascular Development. Curr Tissue Microenviron Rep. 2020;1(3):143–154.

5. Klouda T, Kim Y, Baek SH, et al. Specialized pericyte subtypes in the pulmonary capillaries. Embo j. 2025;44(4):1074–1106.

6. Kim H, Liu Y, Kim J, et al. Pericytes contribute to pulmonary vascular remodeling via HIF2α signaling. EMBO reports. 2024;25(2):616–645-645.

7. Yao Y. Challenges in Pericyte Research: Pericyte-Specific and Subtype-Specific Markers. Transl Stroke Res. 2022;13(6):863–865.

8. Alvino VV, Mohammed KAK, Gu Y, Madeddu P. Approaches for the isolation and long-term expansion of pericytes from human and animal tissues. Front Cardiovasc Med. 2022;9:1095141.

9. Ricard N, Tu L, Le Hiress M, et al. Increased pericyte coverage mediated by endothelial-derived fibroblast growth factor-2 and interleukin-6 is a source of smooth muscle-like cells in pulmonary hypertension. Circulation. 2014;129(15):1586–1597.

10. Mogensen C, Bergner B, Wallner S, et al. Isolation and functional characterization of pericytes derived from hamster skeletal muscle. Acta Physiol (Oxf). 2011;201(4):413–426.

11. Nayak RC, Berman AB, George KL, Eisenbarth GS, King GL. A monoclonal antibody (3G5)-defined ganglioside antigen is expressed on the cell surface of microvascular pericytes. J Exp Med. 1988;167(3):1003–1015.

12. McErlain T, Branco CM, Murgai M. Isolation and Culture of Primary Pericytes from Mouse. Bio Protoc. 2025;15(8):e5288.

13. Yuan K, Orcholski ME, Panaroni C, et al. Activation of the Wnt/planar cell polarity pathway is required for pericyte recruitment during pulmonary angiogenesis. Am J Pathol. 2015;185(1):69–84.

14. James J, Dekan A, Kacar S, et al. Distinct populations of lung capillary endothelial cells and their functional significance. Communications Biology. 2025.

15. Greenwood-Goodwin M, Yang J, Hassanipour M, Larocca D. A novel lineage restricted, pericyte-like cell line isolated from human embryonic stem cells. Sci Rep. 2016;6:24403.

16. Park TI, Feisst V, Brooks AE, et al. Cultured pericytes from human brain show phenotypic and functional differences associated with differential CD90 expression. Sci Rep. 2016;6:26587.

